# Recombination landscape and karyotypic variations revealed by linkage mapping in the grapevine downy mildew pathogen *Plasmopara viticola*

**DOI:** 10.1101/2024.07.03.601951

**Authors:** Etienne Dvorak, Isabelle D. Mazet, Carole Couture, François Delmotte, Marie Foulongne-Oriol

**Affiliations:** INRAE, Bordeaux Sciences Agro, SAVE, ISVV, Villenave d’Ornon, France.; INRAE, MycSA, Villenave d’Ornon, France.

**Keywords:** plant pathogen, genetic map, meiotic recombination, karyotypic anomalies

## Abstract

*Plasmopara viticola*, the causal agent of grapevine downy mildew, is a biotrophic oomycete engaged in atight coevolutionary relationship with its host. Rapid adaptation of the pathogen is favored by annual sexual reproduction which generates genotypic diversity. With the aim of studying the recombination landscape across the *P. viticola* genome, we generated two half-siblings F1 progenies (N=189 and 162). Offspring were genotyped by targeted sequencing of a set of 5,000 informative SNPs. This enabled the construction of unprecedented high-density linkage maps for this species. The reference genome could be assembled into 17 pseudo-chromosomes, anchoring 88% of its physical length. We observed a strong collinearity between parental genomes and extensive synteny with the downy mildew *Peronospora effusa*. In the consensus map, the median recombination rate was 13.8 cM/Mb. The local recombination rate was highly variable along chromosomes and recombination was suppressed in putative centromeric regions. Recombination rate was found negatively correlated with repeats coverage and positively correlated with gene coverage. However, genes encoding secreted proteins and putative effectors were under-represented in highly recombining regions. In both progenies, about 5% of the individuals presented karyotypic anomalies. Aneuploidies and triploidies almost exclusively originated from the male-transmitted chromosomes. Triploids resulted from fertilization by diploid gametes, but also from dispermy. Obligatory sexual reproduction each year may explain the lower level of karyotypic variation in *P. viticola* compared to other oomycetes. The linkage maps will be useful to guide future *de novo* chromosome-scale assemblies of *P. viticola* genomes and to perform forward genetics.

**Summary:** Sexual reproduction contributes to the rapid adaptation of plant pathogens thanks to the recombination of genetic variants. By crossing strains of grapevine downy mildew, we built a set of linkage maps and describe the recombination landscape along the 17 chromosomes of the genome. The recombination rate varied greatly and was higher in gene-rich regions, but not in effector-rich regions. We observed an abnormal number of chromosomes in 5% of the offspring, which was mostly due to the male parent.

## Introduction

Filamentous plant pathogens and their hosts are engaged in coevolution relationships, in which reciprocal selection takes place (Brown and Tellier, 2011). As plants adapt to avoid or limit infection, pathogens counter-adapt to overcome their defenses. This has crucial implications for crop production systems, in which disease control by resistant cultivars can quickly lose efficiency. The evolutionary potential of a plant pathogen depends on its mutation rate, its effective population size, its gene flow capacity and its mode of reproduction (McDonald and Linde, 2002). Among these forces, the prevalence of sexual reproduction plays an important role, because recombination creates original allele associations and leads to an increased genotypic diversity on which natural selection can act. In addition, meiosis itself can generate structural rearrangements and karyotypic variability, which are common in plant pathogens and can have strong impact on life history traits (Mehrabi et al, 2017). In agricultural pathosystems, sexual reproduction can be involved in the emergence of novel pathotypes adapted to resistant cultivars (Ahmed et al, 2012) or different host species (McMullan et al, 2015; Menardo et al, 2016).

Filamentous plant pathogen genomes contain a large number of genes encoding secreted proteins that promote colonization by altering host physiology and suppressing immune responses (Franceschetti et al, 2017). These genes tend to be located in regions rich in repetitive elements. This observation led to the concept of the two-speed genome model, according to which repeat-rich and gene-sparse compartments provide a genomic environment that fosters the fast evolution of genes crucial for host adaptation (Dong et al, 2015). In parallel, fine-mapping of crossing-overs (CO) in fungal pathogens showed that genes involved in host-pathogen interaction are preferentially located in highly recombining regions (Croll et al, 2015; Laurent et al, 2018). This points to a tight link between the functional architecture and the recombination landscape of these genomes. In regions where COs occur frequently, linkage disequilibrium is reduced, which locally increases the efficacy of selection and could therefore boost the speed of pathogen adaptation. Compared to fungal plant pathogen, recombination data remain scarce for oomycetes. Linkage maps have been produced with increasing resolution for several major oomycete pathogens (Hulbert et al, 1988; Van der Lee et al, 1997; Lamour et al, 2012; Matson et al, 2022), but these studies did not focus on recombination per se.

Oomycetes present strong convergence in terms of morphology and lifestyle with true fungi, but they are phylogenetically distant. Their distinctive features include biflagellate zoospores, little to no chitin in cell walls, the absence of an haploid stage and large sexual propagules called oospores. Polyploidy and aneuploidy are frequent in several species, which can complicate genetic analyses (Kasuga et al, 2016; Li et al, 2017). Plant pathogenic oomycetes display a wide range of modes of reproduction, with diverse rates of sexual reproduction (Judelson, 2009). The prevalence of outcrossing also differs between species, as some species are homothallic and reproduce predominantly by selfing (Judelson, 2009). Plant pathogenic oomycete genomes encode hundreds of putative effectors which are often characterized by amino acid sequence motifs such as RXLR/dEER or LXLFLAK (Schornack et al, 2009). They typically occur in large clusters in the genome, suggesting that they evolve by tandem gene duplication (Schornack et al, 2009). It has been suggested that chromosomal rearrangements through unequal COs and/or transposon activity mediate the expansion of RXLR effector families (Matson et al, 2022).

*Plasmopara viticola*, the causal agent of grapevine downy mildew, is a major oomycete pathogen threatening vineyards worldwide (Gessler et al, 2011; Kamoun et al, 2015). It has demonstrated a high ability to adapt by overcoming resistance genes (Wingerter et al, 2021; Paineau et al, 2022) and losing fungicide sensitivity (Chen et al, 2007; Delmas et al, 2017). This high evolutionary potential relies on large population sizes, airborne dispersal and a mixed reproduction system (McDonald and Linde, 2002). At the field level, *P. viticola* populations are panmictic and highly diverse (Gobbin et al, 2006), with a high heterozygosity rate (Dussert et al, 2019). This is favored by outcrossing, as the species is heterothallic with two mating-types denominated P1 and P2 (Wong et al, 2001). Self-incompatibility is controlled by a 570-kb locus exhibiting two highly divergent alleles, with P2 strains being homozygous (MAT-a/MAT-a) and P1 strains heterozygous (MAT-a/MAT-b) (Dussert et al, 2020b). As *P. viticola* is a strictly biotrophic pathogen, sexual reproduction takes place inside the grapevine mesophyll. There, strains of opposite mating-types form gametangia in which meiosis occurs shortly before fertilization. Female oogonia are fertilized by male antheridia and mature into mononucleated oospores, each containing a distinct diploid zygote (Vercesi et al, 1999). Oospores are the only overwintering form under temperate climates. Consequently, the life cycle is characterized by mandatory annual sexual reproduction in the main grape growing areas. Original genotypes are thus generated each year through recombination. A in-depth comprehension of this key driver of the *P. viticola* genome evolution will contribute to a more effective and sustainable management of the disease.

In this study, we assess the relation between meiotic recombination and the genome architecture of an oomycete plant pathogen. We build a set of unprecedented high-density linkage maps based on targeted genotyping-by-sequencing of two large *P. viticola* progenies. We compare the maps obtained for three different parent strains and combine data from the two half-siblings families to obtain a consensus map. Thanks to the linkage data, we generate a pseudo-assembly of the reference genome, making it possible to conduct analyses at the chromosome scale. We compare the local recombination rate along the reconstructed chromosomes with various genomic elements, such as repeats and putative effector genes. We also assess the extent of karyotypic variation in the offspring and identify multiple mechanisms of origin for these anomalies. These results are discussed in light of the evolutionary potential of *P. viticola*.

## Materials and Methods

### Generation of the progenies

As *P. viticola* is an obligate biotrophic organism, sexual reproduction was carried out *in planta* via co-inoculation of strains of opposite mating-types (Fig. 1a). We used single-sporangia strains that were previously characterized by Paineau et al (2022): Pv412 11 (P2) was collected in Switzerland (2010), Pv1419 1 (P1) in Germany (2012), and Pv2543 1 (P1) in Hungary (2013). We generated two progenies with one parent (Pv412 11) in common between the two crosses (Fig. 1b).

**Fig. 1.**
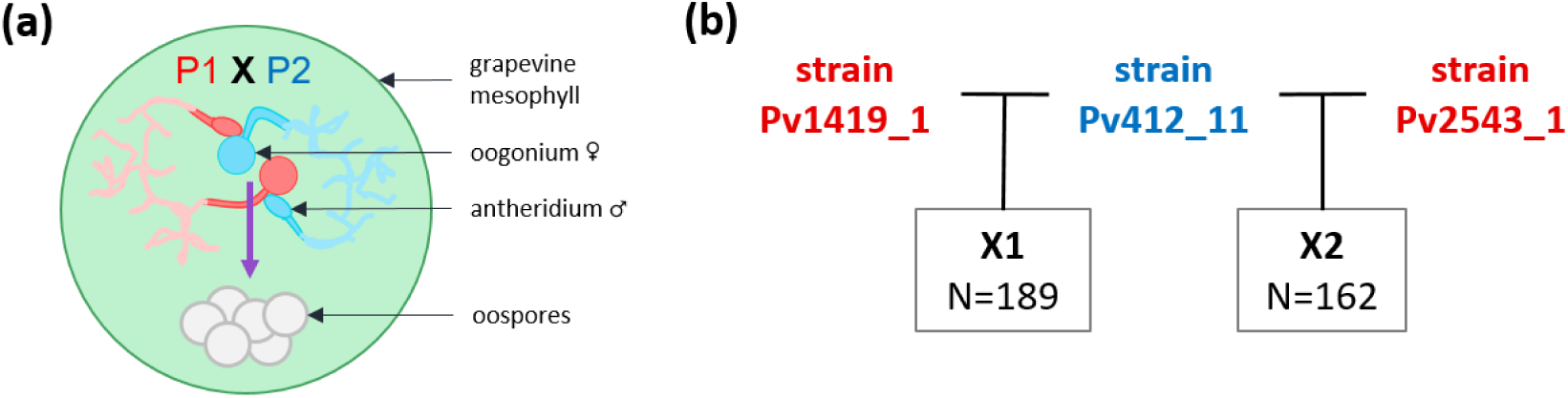
Controlled sexual reproduction of *P. viticola*. **(a)** Gamete-producing organs of opposite mating-types P1 and P2 meeting inside grapevine leaf disc tissues. Fertilized oogonia mature into resting thick-walled oospores. **(b)** Crossing plan carried out in this study. The two crosses have one parent in common, therefore yielding two half-sibling progenies, designated X1 and X2. Mating-types P1 and P2 are respectively represented in red and blue.

All strains used in the study were propagated using leaf tissues sampled from potted grapevine cuttings cv. ’Cabernet-Sauvignon’ grown under greenhouse conditions.

Discs were excised from the seventh leaf from the apex and co-inoculated with the two parents as described in Dussert et al (2020b). Leaf discs in which oospores were observed were put under maturation conditions in the dark at 4°C for 8 to 10 months. Leaf discs were then ground in a glass potter and oospores were retrieved using nylon filters (protocol adapted from Toffolatti et al (2007)). Two steps of filtration were performed to eliminate larger debris (pore size 100 and 60 µm). A third step was done to filter smaller debris while retaining oospores on the filter (pore size 20 µm). The oospores were re-suspended in sterile water with an adjusted concentration of 6-7 per µL, and 15 µL droplets were deposited on 1% water agar plates. These were placed in a growth chamber for 7 to 12 days at 22°C in the dark to trigger germination. The plates were checked daily for macrosporangia formation under a binocular magnifier. Germinating oospores were retrieved one by one with a pipette and inoculated on a disc excised from the fourth leaf from the apex. After successful propagation, a step of single asexual sporangium isolation was performed to ensure that the recovered strains corresponded to a unique genotype, as described in Paineau et al (2022). In total, 189 offspring were obtained for X1, and 162 for X2. (Fig. 1b). Sporulating leaf fragments were desiccated and then stored at -20°C until further use.

### DNA extraction

Strains were multiplied on three detached grapevine leaves (fourth leaf from the apex), and pellets of sporan-giophores and sporangia were collected and treated as described in Dussert et al (2019). DNA was extracted following a CTAB extraction protocol adapted from Möller et al (1992). In particular, protein degradation was performed using 10 µL of Thermo Scientific Proteinase K (20 mg/L, product reference: EO0491). A step of RNA removal was performed after CTAB lysis by adding 10 µL of Qiagen Ribonuclease A (catalog number: 19101) diluted tenfold, followed by incubation at 37°C for 30 minutes. DNA purity and concentration were assessed using respectively a DeNovix DS-11 spectrophotometer and a Qubit 3 fluorometer (Invitrogen).

### SNP Genotyping

Parent strains were sequenced to identify variants suitable for use as genetic markers. Paired-end sequencing was performed at the Genoscope, CEA - Institut de biologie François Jacob (Evry, France) with two runs of Illumina MiSeq 2×250 bp. Following the pipeline used in Paineau et al (2024), variants were called with GATK HaplotypeCaller v4.2.6.1 (McKenna et al, 2010). A total of 3,543,322 potential polymorphic sites were called on the Pv221 reference genome assembly (Dussert et al, 2019).

#### Definition of the set of targeted markers

In order to find high-confidence single-nucleotide polymorphisms (SNPs) that would segregate in the progenies, the vcf file obtained was subjected to a series of filters. Repeated sequences were masked using bedtools v2.27.1 (Quinlan and Hall, 2010), following the annotation available in Dussert et al (2020a). Variants with missing data in any of the parent strains, indels, and multi-allelic SNPs were ignored. After examination of the distribution of GATK annotation values, bi-allelic SNPs were selected using GATK VariantFiltration command with the following parameters: QD *<* 5, QUAL *<* 30, DP *<* 30, SOR *>* 3, FS *>* 40, MQ *<* 50, MQRankSum *<* -5, ReadPosRankSum *<* -5.

To minimize the odds of incorrect calling due to mapping errors, SNPs were considered reliably heterozygous for an individual if minor allele frequency (MAF) *>* 0.4, and homozygous if MAF *<* 0.1. Other SNPs were ignored. After all the steps above, 1,200,172 SNPs were retained.

In our setting, markers were informative when they segregate either in a testcross (1:1) or intercross (1:2:1) configuration. We aimed to define a unique set of markers segregating in the two F1 progenies, which meant SNPs had to be heterozygous in at least one parent for each cross. We preferentially selected markers in the first three types of configurations described in Table 1.

**Table 1.** Informative marker configurations and their expected segregation patterns.

| Genotype configuration | Pv1419_1 | Pv412_11 | Pv2543_1 | segregation in X1 | segregation in X2 |
| --- | --- | --- | --- | --- | --- |
| 1 | H | A | H | 1:1 | 1:1 |
| 2 | A | H | H | 1:1 | 1:2:1 |
| 3 | H | H | A | 1:2:1 | 1:1 |
| 4 | A | H | A | 1:1 | 1:1 |
| 5 | H | H | H | 1:2:1 | 1:2:1 |

A: homozygous ; H: heterozygous

A minimal distance of 6.5 kb between two markers on the same scaffold was chosen. Potential marker positions were then sent to the sequencing facility for the design of specific pairs of probes. Finally, 4996 genomic markers were selected to cover as many scaffolds as possible, to which we added 4 mitochondrial markers.

#### Targeted genotyping-by-sequencing

Parent strains and their offspring were genotyped by targeted sequencing at the INRAE EPGV facility (National Genotyping Center, Evry, France) using Single Primer Enrichment Technology (SPET) (Scaglione et al, 2019) commercialized under the name Allegro. This method is based on a single primer extension reaction to perform multiplex enrichment of a set of thousand of target loci. SPET probes are around 40-bases long and are designed adjacent to a region containing a target variant. This enables the genotyping of the target SNPs but also of additional ones that may surround it. We thereafter refer to these additional variants as flanking variants. Libraries were prepared using the Tecan Allegro Targeted Genotyping V2 kit, with 48 samples pooled per library. One parental genotype was included in all libraries as a control. Sequencing was performed on an Illumina NextSeq 550 (2×150 bp paired-end reads).

#### Calling of targeted SNPs and flanking variants

Reads were mapped on the PV221 reference genome (Dussert et al, 2019) with bwa-mem v2.2.1 (Vasimuddin et al, 2019). Duplicates were removed with the GATK command MarkDuplicatesSpark. Reads with mapping quality *<* 20 were filtered. Probe sequences in the reads were tagged using Tecan Genomics’ tool ProbeFilter (available at https://interval.bio/allegro bioinformatics.html) so that they could be ignored downstream. Variant calling was done using GATK HaplotypeCaller in GVCF mode. Then, GVCFs were consolidated scaffold by scaffold using GATK GenomicsDBImport, before joint calling of genotypes with GATK GenotypeGVCFs. Finally, all files were merged using GATK GatherVcfs. Variants with read depth (DP) ¡ 10 were set to missing data. Additionally, they were considered heterozygous only if MAF *>* 0.1.

#### Mitochondrial heredity

In oomycetes, mitochondria are solely transmitted by female oogonia that provide the cytoplasm of the future oospores (Martin, 1989; Judelson, 2009). Four mitochondrial markers were used to follow this heredity and determine the female parent of each offspring.

### Linkage maps construction

As *P. viticola* is a highly heterozygous self-incompatible organism (Wong et al, 2001; Dussert et al, 2019), we used a two-way pseudo-testcross mapping strategy. Heterozygous markers in each parent were used to build distinct parental maps, with each parent acting as a ”tester” for the other (Hulbert et al, 1988; Grattapaglia and Sederoff, 1994).

#### Parental maps

Parental linkage maps were independently built using r/ASMap v1.0.4 (Taylor and Butler, 2017) in successive steps. Individuals with excessive heterozygosity rates (*>* 0.6) were ignored, as this could be due to genotype mixture or polyploidy. The presence of identical genotypes was checked (*>*95% marker identity). The marker phase in each parent was initially unknown, so all markers were added twice, once for each opposite phase, as mentioned in Brelsford et al (2016). Duplicate linkage groups (LGs) were subsequently removed. Markers were pulled out when more than 20% of the genotypes were missing or if their segregation pattern was significantly distorted (Chi-squared test, p *<* 0.1 / number of markers).

First of all, framework maps were constructed with only testcross target markers (homozygous in one parent and heterozygous in the other, as described in Table 1). Testcross flanking markers were added to each map if they belonged to scaffolds with little or no markers already positioned. A few intercross (heterozygous in both parents) target markers were incorporated to densify the maps where long gaps remained. For these, 0/1 genotypes were set as missing because of the impossibility of knowing the parental origin of the alleles. Consequently, the missing data threshold was bypassed for these markers. Two LGs of the X1 maps exhibited large gaps because of the removal of closely linked markers that were moderately distorted. We decided to put back these groups of markers as they allowed to complete the maps without aberrant behavior (see Results).

One marker per unique segregation pattern was retained for map construction with the mstmap function. LGs were separated using a LOD threshold of 8. Genetic distances were calculated with the Kosambi mapping function. After the initial marker ordering, distances were recomputed by applying a genotyping error rate of 0.5%, in accordance with the repeatability of the control samples. Then, the maps were manually curated by using the compareorder function. When contradictions were observed between genetic and physical orders, markers were switched if doing so increased likelihood and reduced LG length. Finally, co-segregating markers were added back.

#### Consensus linkage map

Scaffolds were ordered and oriented by combining information from the four final parental maps with ALLMAPS (Tang et al, 2015). As there were very few discrepancies between genetic and physical order (Spearman’s *ρ* ranging from 0.98 to 1.00), markers from all maps were ordered according to their physical position in the reference genome. In order to integrate linkage information from both families, the consensus map was built using Lep-MAP3 (version available on 20-08-2023) (Rastas, 2017). To avoid the spurious estimation of recombination fraction between markers flanking the same sequenced target, only the most informative ones were kept for each targeted region (i.e. markers for which there were as many heterozygous parents as possible). The genotypes of the parents and the two half-sib progenies were determined using the ParentCall2 command with option halfSibs=1. Markers were phased and the genetic distances were calculated for each LG with the Kosambi mapping function and a genotyping error rate of 0.5%. This was achieved using the OrderMarkers2 command with options improveOrder=0, useKosambi=1, minError=0.005, sexAveraged=1, outputPhasedData=1.

### Genomic features

#### Synteny

The synteny between the anchored *P. viticola* scaffolds and the chromosome-level assembly of *Pe. effusa* was assessed using SynMap2 (Haug-Baltzell et al, 2017). Coding sequences positions were retrieved from the published annotations (Dussert et al, 2019; Fletcher et al, 2022) and were aligned with the ”LAST” algorithm (default parameters).

#### Centromere identification

Putative centromeric regions were identified by the presence (BLASTn alignment length *>* 500 bp) of a*Phytophthora sojae* Copia-like Transposon proposed as a distinctive feature of oomycete centromeres (Fang et al, 2020). The regions were confirmed via synteny with the *Pe. effusa* centromeres described by Fletcher et al (2022).

#### Pseudo-assembly and annotations

Thanks to the linkage data, the scaffolds of the reference genome were joined and oriented using ALLMAPS, adding 100 Ns between each one. Annotations from Dussert et al (2019) were lifted over to this pseudo-assembly, including the positions of predicted secreted protein genes and putative effector genes.

#### Estimation of the recombination rate

Using genetic distances from the consensus map, the recombination rate in cM/Mb was approximated in the pseudo-assembly using Locally Estimated Scatterplot Smoothing (LOESS) implemented in r/MareyMap v1.3.6 (Rezvoy et al, 2007), with parameters span=0.1 and degree=2. The estimation was retrieved every 10 kb for graphical representation and statistical analyses.

#### Gene enrichment analysis

Ten-kb genomic bins for which recombination rate could be estimated were regrouped by deciles. Limits were calculated using the quantile() function implemented in R v1.4.2. Each decile was tested for enrichment or depletion in secreted protein and putative effector genes (hypergeometric test, *α* = 0.05).

### Identification of karyotypic anomalies

Aneuploid and triploid individuals were first identified by their highly excessive number of apparent COs inone or several LGs of a parental map. For each individual, the number of detected cross-overs in each parental LG was obtained using the countXO function of r/qtl v1.60 (Arends et al, 2010). These observations were supported by the examination of allelic ratios in relevant LGs or the entire genome. Aneuploidies and partial duplication/deletions could be additionally corroborated by comparing reads mapping coverage between LGs. The affected parental linkage map designated from which parent the anomaly originated. The sequencing depth of the targeted SNPs was retrieved from the reads aligned to the reference genome using mosdepth v.0.2.5 (Pedersen and Quinlan, 2018). For partial duplication/deletion and copy-neutral losses of heterozygosity, the estimated boundaries correspond to the region with continuous switching of markers from one phase to another (as mentioned in Lamour et al (2012)). Such a switch indicates that a portion of the chromosome is in fact fully homozygous or fully heterozygous for the considered markers.

Aneuploid strains were kept for the construction of linkage maps as they still provided valuable information for most of the genome. The false recombination events in affected LGs had little impact thanks to the allowance for 0.5% of genotyping errors. Triploids were all put aside previously because of their excessive heterozygosity, as mentioned above.

### Assessment of origin of triploidy

To identify the most likely mechanism of origin of triploidy for each individual, we analyzed the allelic composition of the two sets of chromosomes transmitted by only one parent. Specifically, we determined if parental heterozygosity was maintained (non-reduction) or reduced to homozygosity in pericentromeric markers, as they do not recombine. By combining observations on multiple chromosomes, we could consider and rule out certain mechanisms for each individual, as explained in Zaragoza et al (2000).

Failure of disjunction in meiosis I is characterized by non-reduced pericentromeric markers, failure of disjunction in meiosis II by reduced markers, and fertilization by two different gametes leads to a random mixture of both. Consequently, we reconstructed the alleles transmitted by each parent for pericentromeric markers in the LGs in which putative centromeres were confidently identified. The number of copies of each allele was estimated using the allelic ratios of the markers. For example, let us consider a marker for which parent 1 (genotype AA) passed on one copy, and parent 2 (AB) two copies. If the observed genotype of the offspring is AA, then parent 2 necessarily transmitted two ”A” copies. When the observed genotype is AB, there are two possibilities: 1) if the proportion of ”A” is statistically equal to 2/3, we can infer that parent 2 transmitted one ”A” copy and one ”B” copy ; 2) if this proportion corresponds instead to 1/3, then it means that two ”B” copies were passed on. The proportions were tested using a chi-squared test with *α* = 0.05.

## Results

### Targeted Genotyping-by-Sequencing of informative SNPs

The progenies were genotyped by targeted sequencing of informative nuclear markers. Out of the 4,996 targeted regions, 4,990 were successfully sequenced, but two markers appeared tri-allelic and were therefore ignored. The mean sequencing depth of the targets was 173x (min. 51x, max. 506x). The repeatability of the genotyping of target SNPs in the control samples was 99.4%. Individuals were successfully genotyped for 97% of all SNPs on average (min. 91%). The Allegro technology provides sequences up to a few hundred bases upstream and downstream of the target SNPs. Thus, a total of 75,812 potentially informative flanking variants were called in X1, and 41,215 in X2. Overall, 4,988 high-confidence target markers were available for the construction of the linkage maps, to which 781 flanking variants were added to reduce gaps.

### Mitochondria are uniparentally inherited

Beside nuclear SNPs, we followed the inheritance of four mitochondrial markers. As expected, mitochondria were uniparentally inherited. Both parents in each cross could transmit it, demonstrating that each strain produced male and female gametes. However, the sex contributions were uneven in X1 (Pearson’s chi-squared test, p=1.2e-4), in which Pv1419 1 was the female parent for two thirds (64.0%) of the offspring. No such imbalance was observed in X2 (p=0.43).

### Construction of a set of parental and consensus linkage maps

#### Markers are consistently gathered into 17 linkage groups

One linkage map for each parent of each cross was built using primarily testcross target markers (segregating 1:1). Between 1,405 and 1,894 markers were positioned depending on the parent strain considered (Table 2). Each map is composed of 17 LGs (Table 2), which correspond in all likelihood to the number of chromosome pairs. The lengths of the two Pv412 11 maps were similar (1167.5 cM in X1 versus 1221.7 cM in X2), and a bit shorter than the Pv2543 1 map (1379.2 cM). The Pv1419 1 map stands out as much longer (2073.4 cM), with a higher number of unique segregation patterns (946 versus 717 to 842) (Table 2). The average spacing between markers ranged from 0.7 cM in Pv412 11-X2 to 1.1 cM in Pv1419 1-X1. Some moderate gaps remained, but they were located on different LGs depending on the map (see Figure S1 for other LGs). For example, the largest gap in Pv412 11-X2 (11.4 cM) was located in LG6, while it was found in LG12 in the Pv1419 1-X1 map (18.8 cM) (Table 2).

**Table 2.** Features of the four parental linkage maps and the consensus map.

| map | Pv412.11<br>X1 | Pv412.11<br>X2 | Pv1419.1<br>X1 | Pv2543.1<br>X2 | consensus |
| --- | --- | --- | --- | --- | --- |
| linkage groups | 17 | 17 | 17 | 17 | 17 |
| markers | 1405 | 1894 | 1879 | 1578 | 4509 |
| markers with unique segregation pattern | 717 | 842 | 946 | 728 | 2739 |
| length (cM) | 1167.5 | 1221.7 | 2073.4 | 1379.2 | 1473 |
| average spacing (cM) | 0.8 | 0.7 | 1.1 | 0.9 | 0.3 |
| maximum spacing (cM) | 13.4 | 11.4 | 18.8 | 17.3 | 5.8 |
| anchored scaffolds | 144 | 142 | 143 | 120 | 154 |
| Total anchored length (Mb) | 80.13<br>(86.2%) | 79.93<br>(86.0%) | 80.33<br>(86.4%) | 76.19<br>(82.0%) | 81.64<br>(87.8%) |

#### The parental genomes are highly collinear

The parental maps were remarkably collinear between them and with the reference genome (correlation of genetic and physical order *ρ >*= 0.98) (Fig. 2). No large-scale structural variations were observed at the resolution level of the linkage maps.

**Fig. 2.**
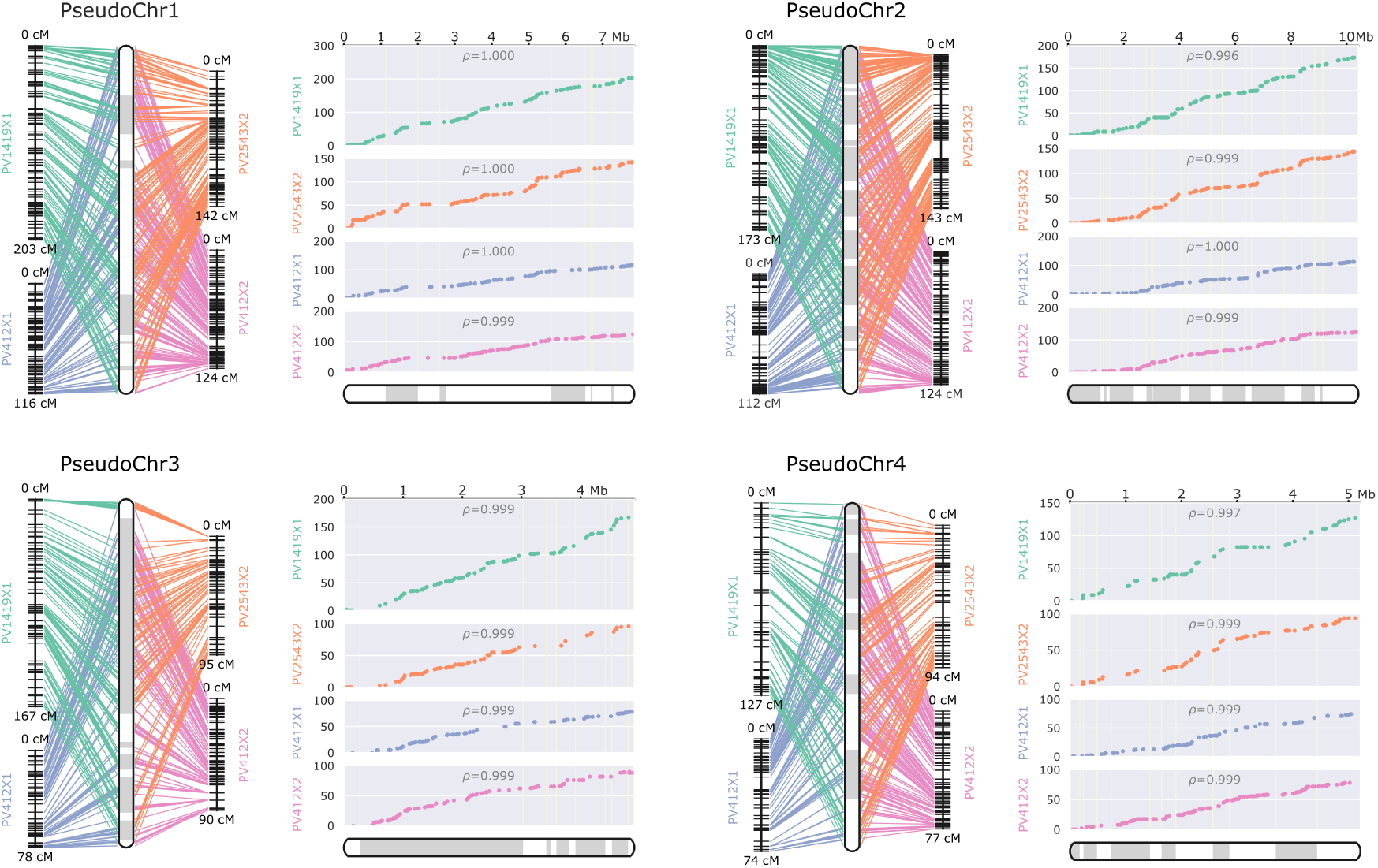
*P. viticola* parental linkage maps. The first four linkage groups are shown (see Fig. S1 for the others). The scaffolds of the reference assembly were anchored and oriented thanks to the linkage information from the four parental maps. They were joined together as pseudochromosomes, which are represented at the center of the left panel and the bottom of the right panel. The scaffolds’ limits are signaled by the switch between white and gray sections. **Left panels:** correspondence between the physical positions and the genetic positions in the linkage groups of the four parental maps. **Right panels:** Genetic positions (y-axis) versus physical positions (x-axis). A strong slope implies a highly recombining region, whereas a rather flat line indicates little or no recombination in the interval. The Spearman’s *ρ* measures the rank correlation between the genetic and the physical order for each LG. Plots were generated using ALLMAPS.

#### The recombination activity varies between strains

The recombination patterns appear similar between all parental maps as can be seen in the right panels of Fig. 2. Interestingly, much more recombination events were detected in Pv1419 1 meiosis than in other parents (from 51% to 80% more COs on average) (Fig. S2). This is reflected by its inflated linkage map length and a higher average spacing between markers (Table 2). The increase in recombination was evenly spread across all LGs (Fig. 2).

The number of COs between maternal and paternal meiosis slightly differed for all parents, although not always in the same direction. More COs occurred in female meiosis for Pv412 11 and Pv2543 1 (+8.0 to 15.6% compared to male meiosis, p*<*0.034), but the opposite was observed for Pv1419 1 (-10.4%, p=0.0018) (Fig. S2).

#### Two regions exhibit distortion of segregation in one cross

Two large regions on different LGs showed a skewed segregation in X1. Markers segregated approximately at a 1:2 frequency instead of the expected 1:1 ratio. Each region was actually affected in only one parental map and the corresponding intervals were free from distortion in the other map. They were located on LG2 in the Pv1419 1 map, and LG13 on the Pv412 11 map. The corresponding physical segments stretched over 0.94 and 1.99 Mb respectively. Notably, the LG13 region was not distorted in the Pv412 11 map obtained inthe other cross. Thus, the segregation distortions appear to result from an incompatibility between a specific combination of alleles at two different loci in X1.

#### The consensus linkage map covers most of the genome length

We took advantage of the high collinearity between the parental genomes to build a consensus linkage map integrating the genotyping data of the two offspring. As desired initially, most markers (97.6%) were informative in both crosses, ensuring a good accuracy in the computing of genetic distances. This resulted in the positioning of 4,509 markers (2,739 unique positions) (Fig. 3), with a reduced spacing between markers and much shorter gaps. In total, 154 scaffolds representing 87.8% of the genome length were anchored in a single map (Table 2). The rest of the genome consists of small repeat-rich scaffolds on which SPET probes were difficult to design.

**Fig. 3.**
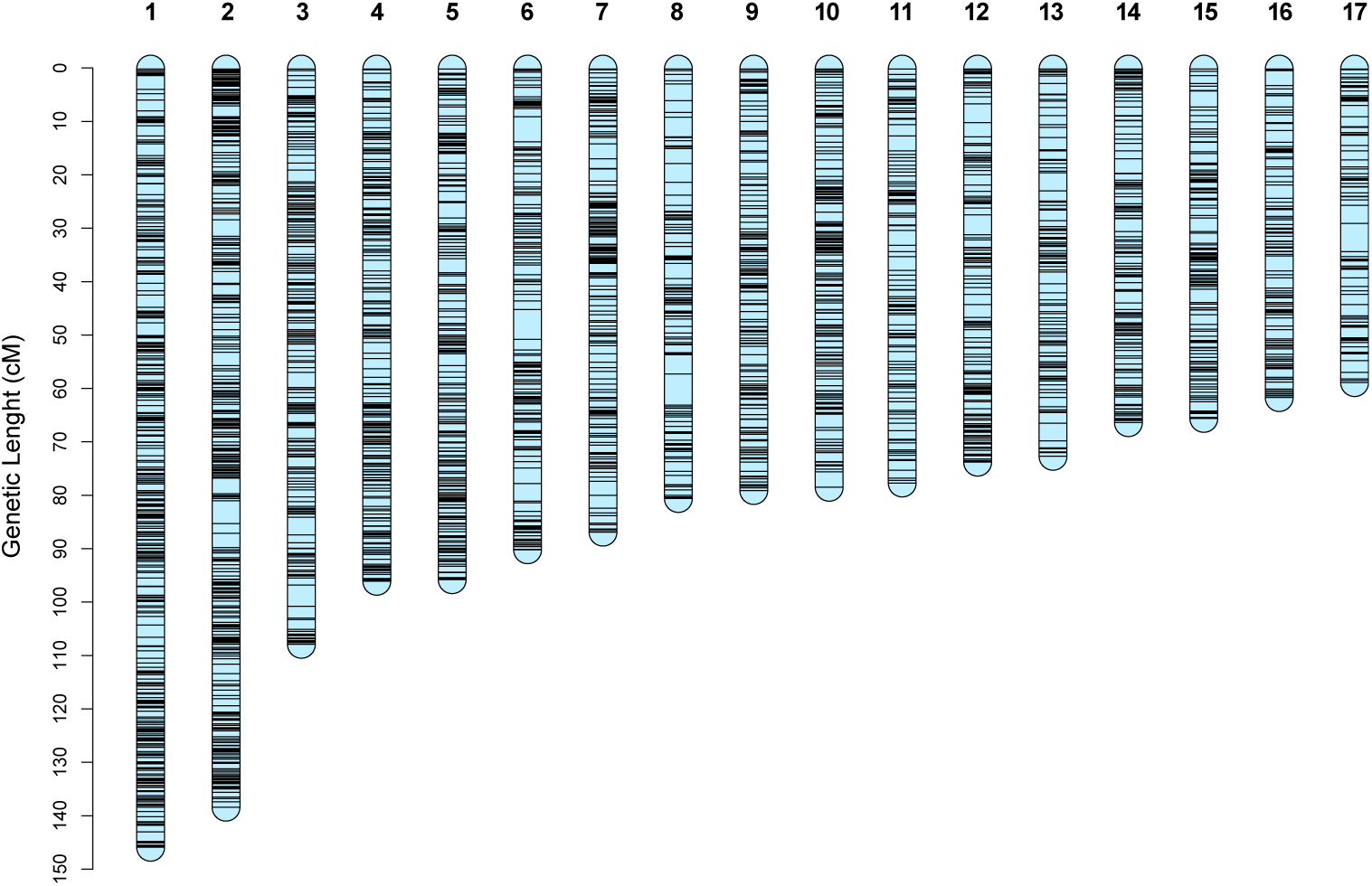
Genetic length and marker distribution of the 17 linkage groups in the consensus linkage map of *P. viticola*. The scale on the left indicates the genetic distances in centiMorgan determined by the Kosambi mapping function. Plot made using r/LinkageMapView v2.1.2.

The LGs were numbered from the longest to the shortest according to their genetic length in this consensus map.

#### The linkage groups are highly syntenic with the Peronospora effusa chromosomes

The 154 anchored *P. viticola* scaffolds were aligned with the chromosome-scale assembly of *Pe. effusa*, the spinach downy mildew pathogen (Fig. 4). Coding sequences from scaffolds in the same LG mapped consistently to the same *Pe. effusa* chromosome, indicating a high level of synteny between the two genomes.

**Fig. 4.**
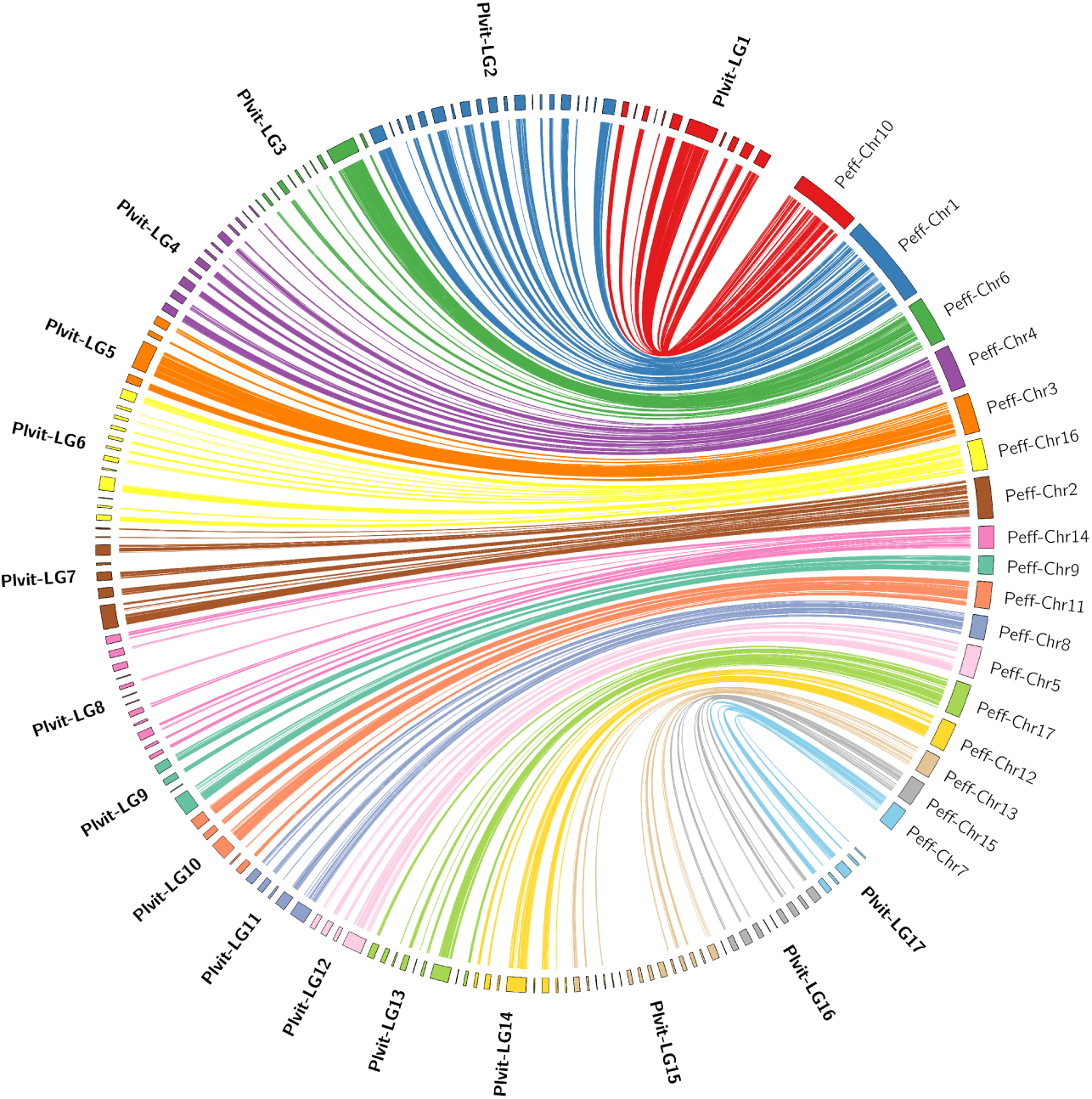
Synteny between the grapevine and spinach downy mildew genomes. On the left, *P. viticola* scaffolds are regrouped and oriented according to the linkage maps, and correspond to 87.8% of the genome length (Table 2). On the right are represented the 17 *Pe. effusa* chromosomes. Each line joins the positions of matching coding sequence (6365 CDS). Non-anchored scaffolds (not shown) did not display synteny with multiple *Pe. effusa* chromosomes. The plot was made using circos v.0.69.9 (Krzywinski et al, 2009)

### Pseudo-assembly of the genome enables the study of the recombination landscape

Thanks to the linkage data, scaffolds of the reference reference could be assembled into 17 pseudochromosomes.

#### The recombination landscape is non-uniform

The genetic length of the LGs and the physical length of the corresponding pseudochromosomes are highly correlated (r=0.88 for the consensus map, p*<*2.2e-16). Using the consensus map, the recombination rate was estimated in 10-kb bins. It fluctuated importantly along all chromosomes (Fig. 5), from 0.0127 cM/Mb on average in the 10% least recombining bins to 63.5 cM/Mb on average for the top 10% (Fig. 6a). Marker density was calculated every 10 kb in 100-kb sliding windows and no correlation was found with the recombination rate (*ρ*=-0.03). In all 10-kb bins, the percentage of bases covered by genes and repeats was calculated. The recombination rate was positively correlated with gene coverage (*ρ*=0.32, p*<*2.2e-16), while the correlation with repeat coverage was negative (*ρ*=-0.39, p*<*2.2e-16). These trends were also visible in the comparison between deciles of recombination rate (Fig. 6a).

**Fig. 5.**
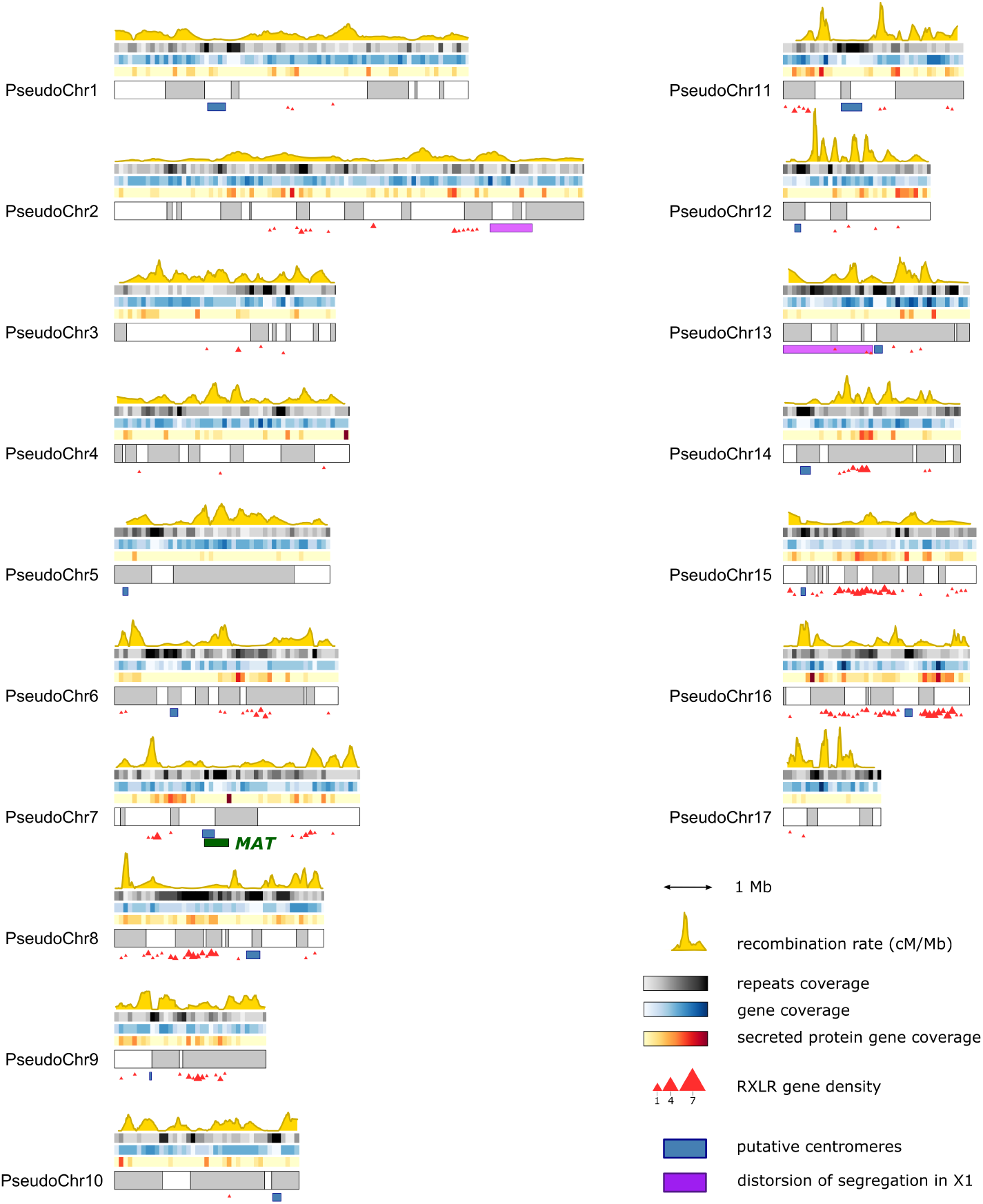
Recombination landscape and genomic features along the reconstructed pseudochromosomes of *P. viticola*. The switch between gray and white sections signals the alternation between scaffolds. On top of the pseudochromosomes is shown, from the top to the bottom: the fraction of bases covered by repeats, genes, and secreted protein genes (nt/kb), all calculated on 100-kb windows. The yellow curve represents the recombination rate in cM/Mb, estimated from the consensus linkage map. Under the pseudochromosomes are displayed notable genomic regions: putative centromeres (in blue), regions affected by segregation distortion in X1 (in purple) and the mating-type locus overlapping with the centromeric region of the pseudochromosome 7 (in green). The density of RXLR genes (as defined in Dussert et al (2019)) is represented by red triangles.

**Fig. 6.**
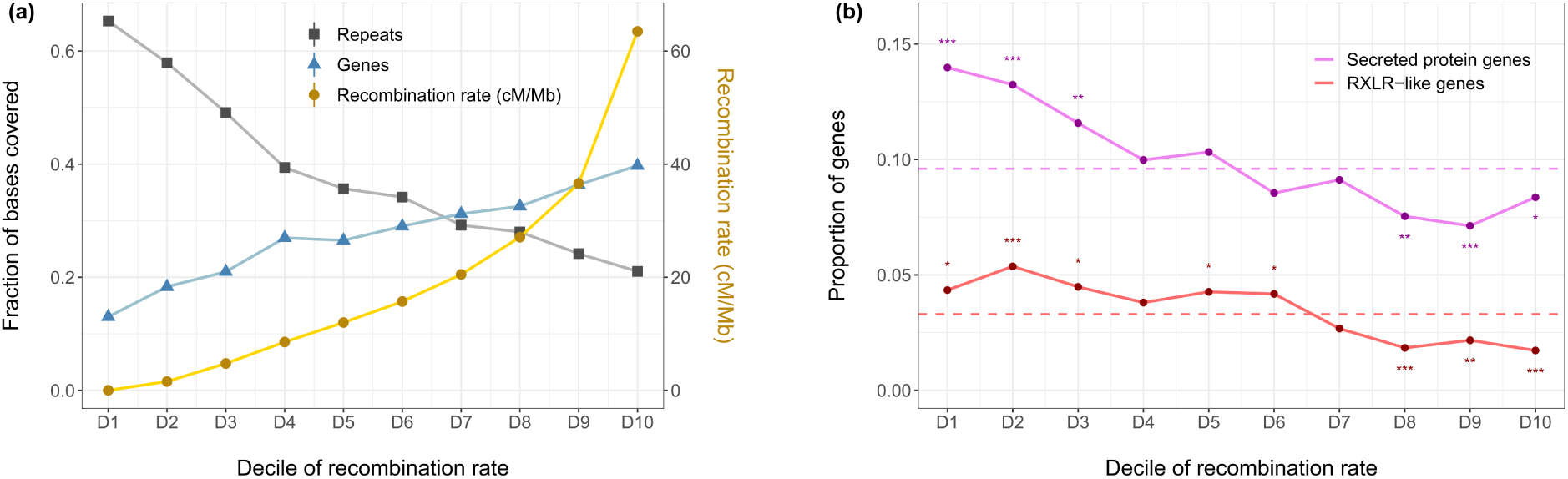
(a) Coverage by genes and repeats in each decile of recombination rate. X-axis : 10-kb genomic bins grouped by deciles of increasing recombination rate in the consensus *P. viticola* linkage map. Right Y-axis : average recombination rate (cM/Mb) in each decile. Left Y-axis : Fraction of bases covered by repeated or coding sequences in each decile. **(b) Enrichment and depletion of secreted protein and RXLR-like genes in each decile of recombination rate.** Dashed lines signal the genome-wide proportion of the genes of interest. Asterisks signal significant enrichment or depletion in each decile (hypergeometric test, ”*” : p ¡ 0.05, ”**” : p ¡ 0.01, ”***” : p ¡ 0.001).

Lack of recombination could be observed in several segments on most pseudochromosomes (Fig. 5). Some of them were likely to be centromeric regions. Based on the presence of specific repeated sequences, putative centromeres were confidently identified in 13 pseudochromosomes out of 17. As expected, they clearly corresponded to repeat-rich and gene-poor regions. (Fig. 5).

#### Putative effectors are relatively depleted in highly recombinant regions

RXLR-like genes were mostly organized in clusters and particularly abundant in some pseudochromosomes, such as PseudoChr8, 15 and 16 (Fig. 5). By contrast, they were absent or very scarce in half of the pseudochromosomes. The positive correlation between gene content and recombination rate at the 10-kb scale disappeared when considering only secreted protein or RXLR-like genes (*ρ*=0.03 and -0.04 respectively). In fact, both secreted protein and RXLR-like genes were significantly more abundant than expected in the three less recombining deciles of the genome, and significantly depleted in the three most recombining ones (hypergeometric test, *α* = 0.05) (Fig. 6b).

#### The mating-type locus is pericentromeric

The 570-kb MAT locus overlaps with the centromere of the pseudochromosome 7 (Fig. 5). The entire locus showed little to no recombination, with different behaviors depending on the mating-type. In the P2 parent Pv412 11 (MAT-a/MAT-a), two COs were detected in the offspring. By contrast, no recombination took place in P1 parents (MAT-a/MAT-b) that possess two divergent alleles at the locus.

### Karyotypic anomalies are found in both progenies

#### Aneuploidies and triploidy originate quasi-exclusively from male gametes

Atypical karyotypes were identified by combining read depth data and allelic ratios for the markers (Fig. 7). These data were consistent with the abnormal phase switches observed in the parental linkage maps. Both progenies exhibited similar levels of aneuploid individuals (1.9% in X1, 2.1% in X2) and triploids (3.2% in X1, 4.3% in X2). Aneuploidies affected various LGs, but their limited number made it difficult to draw conclusions on the frequency at which each LG was concerned (Table 3). In diploids, trisomic individuals were the most common, but one monosomic strain was identified as well (Fig. S3). Some triploids in X2 displayed additional aneuploidies, as exemplified in Fig.7b. Moreover, partial deletion or duplication of chromosome ends were also detected and spanned large segments (*>*1 Mb) (Fig. S3).

**Fig. 7.**
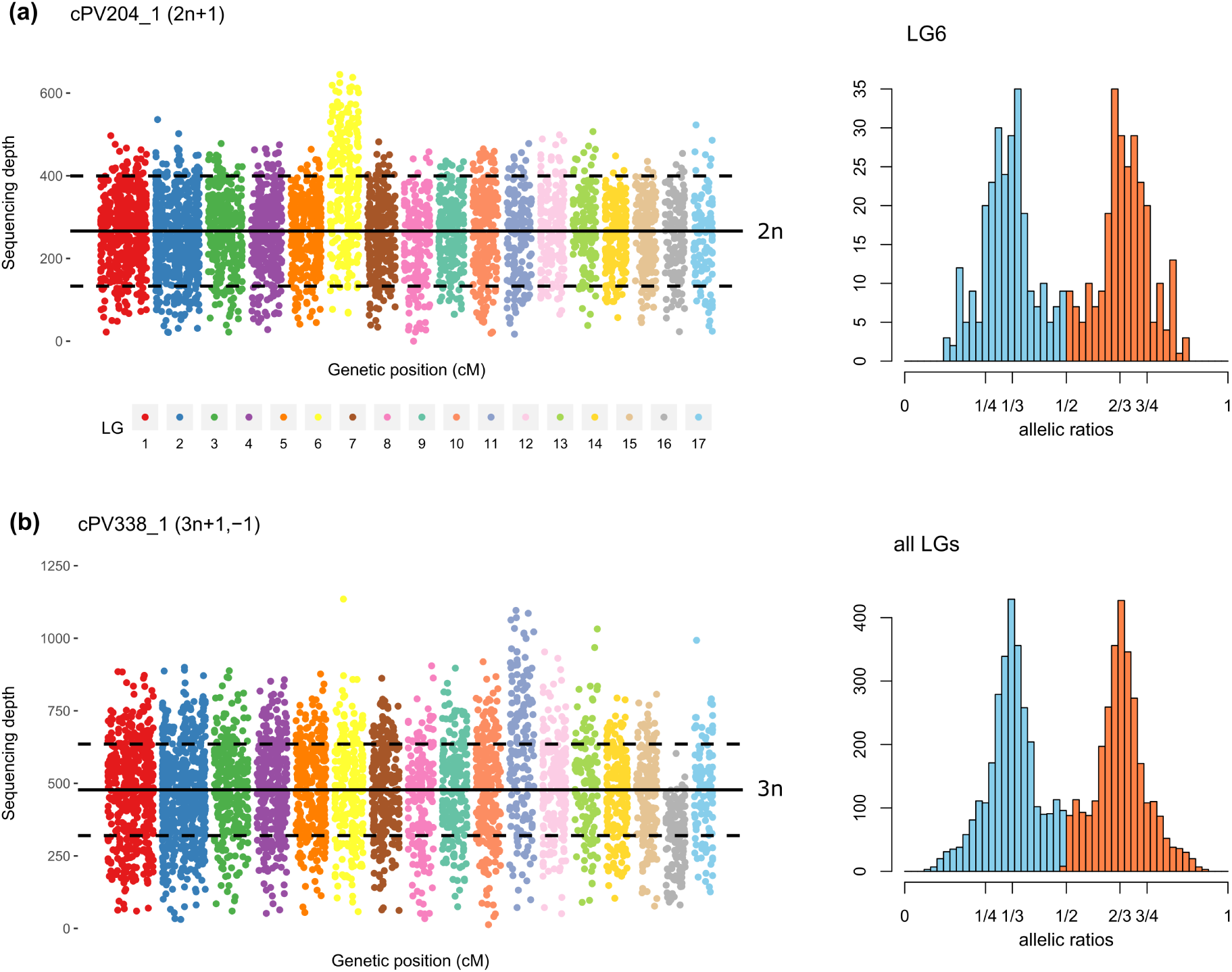
Examples of karyotypic anomalies in *P. viticola* offspring. **(a)** On the left is represented the read depth of all markers across all linkage groups for cPV204 1, a trisomic individual carrying an extra copy of LG6. The full line represents the average sequencing depth, and the dashed lines are plotted at +50% and -50% of this value. On the right, allelic ratios of heterozygous markers for LG6 show peaks at 1/3 and 2/3 (minor allele in blue, major allele in red). **(b)** Same data for cPV338 11, a triploid individual displaying additional aneuploidies: 4 copies of LG11 (tetrasomy) and 2 copies of LG16 (disomy). On the left panel, dashed lines are plotted at +33% and -33% of the average depth. On the right panel, allelic ratios of heterozygous markers across the entire genome are consistent with a global ploidy number of 3. Additional examples are available in Figure S3.

**Table 3.** Diploid offspring affected by karyotypic anomalies.

| cross | strain | origin of the anomaly | male parent | anomaly | affected linkage group | approximate length |
| --- | --- | --- | --- | --- | --- | --- |
| X1 | cPV57_1 | Pv412_11 | Pv412_11 | trisomy | 15 | - |
| X1 | cPV101_1 | Pv412_11 | Pv412_11 | partial deletion | 8 | 1 Mb |
| X1 | cPV138_1 | Pv412_11 | Pv412_11 | trisomy | 16 | - |
| X1 | cPV204_1 | Pv412_11 | Pv412_11 | trisomy | 6 | - |
| X1 | cPV236_1 | Pv412_11 | Pv412_11 | trisomy | 17 | - |
| X1 | cPV243_1 | Pv412_11 | Pv412_11 | partial duplication | 2 | 1 Mb |
| X2 | cPV308_1 | Pv412_11 | Pv412_11 | trisomy | 17 | - |
| X2 | cPV329_1 | Pv412_11 | Pv412_11 | partial deletion | 11 | 1.4 Mb |
| X2 | cPV409_1 | Pv412_11 | Pv412_11 | trisomy | 3 | - |
| X2 | cPV419_1 | Pv412_11 | Pv412_11 | partial deletion | 8 | 1.5 Mb |
| X2 | cPV462_1 | Pv412_11 | Pv412_11 | monosomy | 16 | - |

Strikingly, all these anomalies were almost systematically transmitted via the male parent (Table 3 and 4). There was only one instance of an extra chromosome inherited from the female gamete (LG12 of the triploid cPV479 1). Notably, simple aneuploids in both progenies resulted exclusively from unbalanced male gametes produced by the common parent Pv412 11. All partial deletions and duplications also originated from this strain.

#### Multiple mechanisms, including dispermy, lead to triploidy

All parent strains were found to occasionally transmit an extra set of chromosomes via male gametes, resulting in triploid offspring (Table 4). To elucidate the mechanism causing this phenomenon, we took advantage of the information available from pericentromeric markers. Thus, we deduced which alleles were transmitted by the male parent for each of the 13 LGs in which a putative centromere was reliably identified (Fig. 5). Briefly, the pattern of reduction or non-reduction to homozygosity of these markers informed us on the most likely origin of the two male chromosome sets (Table 4).

**Table 4.** Triploid offspring and their diverse mechanisms of origin.

| cross | strain | origin of the extra chromosome set | male parent | additional aneuploidy* | reduced pericentromeric markers | non-reduced pericentromeric markers | most likely origin of triploidy |
| --- | --- | --- | --- | --- | --- | --- | --- |
| X1 | cPV33.1 | Pv412.11 | Pv412.11 |  | 77% | 23% | dispermy |
| X1 | cPV34.1 | Pv1419.1 | Pv1419.1 |  | 42% | 58% | dispermy |
| X1 | cPV74.1 | Pv412.11 | Pv412.11 |  | 0% | 100% | non-disjunction in meiosis I |
| X1 | cPV79.1 | Pv412.11 | Pv412.11 |  | 69% | 31% | dispermy |
| X1 | cPV89.1 | Pv412.11 | Pv412.11 |  | 38% | 62% | dispermy |
| X1 | cPV213.1 | Pv412.11 | Pv412.11 |  | 100% | 0% | non-disjunction in meiosis II |
| X2 | cPV305.1 | Pv412.11 | Pv412.11 |  | 100% | 0% | non-disjunction in meiosis II |
| X2 | cPV309.1 | Pv412.11 | Pv412.11 | tetrasomy LG14 | 100% | 0% | non-disjunction in meiosis II |
| X2 | cPV313.1 | Pv412.11 | Pv412.11 |  | 46% | 54% | dispermy |
| X2 | cPV338.1 | Pv2543.1 | Pv2543.1 | tetrasomy LG11 ; disomy LG16 | 100% | 0% | non-disjunction in meiosis II |
| X2 | cPV447.1 | Pv2543.1 | Pv2543.1 |  | 50% | 50% | dispermy |
| X2 | cPV449.1 | Pv2543.1 | Pv2543.1 |  | 33% | 67% | dispermy |
| X2 | cPV479.1 | Pv2543.1 | Pv2543.1 | tetrasomy LG12 and LG17 | 100% | 0% | non-disjunction in meiosis II |

\*All additional aneuploidies originated from male gametes as well, except in the case of LG12 in cPV479.1, for which both parents transmitted two chromosome copies.

In both crosses, we identified several mechanisms that led to a triploid zygote. In half of all cases, non-disjunction of chromosomes during meiosis led to the production of diploid male gametes. These non-disjunction events occurred over the first meiotic division (consistently non-reduced markers, N=1) or during the second one (consistently reduced markers, N=5) (Table 4). Interestingly, we also observed triploids carrying a mix of reduced and non-reduced markers, which can only be explained by dispermy, i.e. fertilization by two independent haploid male gametes (N=7). Notably, this last mechanism involved all parent strains (Table 4). Therefore, triploidy was caused by abnormal meiosis as well as atypical fertilization.

Finally, some triploids displayed additional aneuploidies in X2. By checking the allelic ratio of variants homozygous in both parents, we found that the female gamete likely transmitted a balanced number of chromosomes in all but one case mentioned in the paragraph above (Table 4).

#### Some individuals present copy-neutral Loss Of Heterozygosity

In total, four individuals displayed long runs of marker homozygosity without evidence of partial deletion ofthe chromosome (Table 5). This Loss Of Heterozygosity (LOH) likely originated from the conversion of the chromosome segment to the other haplotype present in its homologous counterpart, which necessarily occurred post-fertilization. Two strains presented LOH in almost the entire LG15. The interval corresponds to the entire long arm of the chromosome, which is acrocentric according to the putative centromere position (Fig. 5).

**Table 5.** Offspring showing loss of heterozygosity on chromosomal segments.

| cross | strain | lost haplotype | affected link-<br>age group | approximate length |
| --- | --- | --- | --- | --- |
| X1 | cPV108_1 | Pv412_11 | 12 | 320 kb |
| X1 | cPV222_1 | Pv1419_1 | 7 | 730 kb |
| X2 | cPV394_1 | Pv412_11 | 15 | 3.7 Mb |
| X2 | cPV457_1 | Pv412_11 | 15 | 3.7 Mb |

## Discussion

We established a high-density consensus linkage map of the *P. viticola* genome, allowing us to study recombination activity and its variation across the genome. Thanks to the linkage data, we generated a pseudo-assembly of the reference genome, making it possible to conduct analyses at the chromosome scale. This work provides valuable insights on the relation between recombination and architecture of oomycete genomes. To our knowledge, it is also the first unequivocal description of polyploidy in a downy mildew pathogen.

### Chromosome-scale pseudo-assembly

Obtaining contiguous assemblies for highly heterozygous oomycetes remains challenging. For such species, linkage maps can still bring decisive inputs. Based on the identified LGs, scaffolds were assembled into 17 chromosomes, which correspond to the ancestral number of chromosome pairs of downy mildews (Fletcher et al, 2022). In total, about 88% of the *P. viticola* genome length could be anchored. For comparison, 89% of the *Ph. infestans* genome was assembled into chromosomes using similar linkage data in addition to an optical map (Matson et al, 2022). In our case, unplaced sequences are small repeat-rich scaffolds on which markers were difficult to define in our setting. Some of them probably correspond to centromeric regions, as we could not reliably position the centromeres in 4 out of 17 chromosomes. Interestingly, we observed a seemingly perfect macro-synteny with the telomere-to-telomere assembly of *Pe. effusa*, which was also noted between other recent highly contiguous downy mildew genomes (Fletcher and Michelmore, 2023). The order of scaffolds determined using the linkage map was congruent with the order inferred by synteny. Thus, high-quality reference genome assemblies could be used to guide assemblies of non-model downy mildew species by taking advantage of the high level of synteny between Peronosporaceae (Seixas et al, 2021).

### Collinearity and compatibility between parental genomes

The genomes of the three parent strains are highly collinear, indicating that structural variations are probably limited in European populations of *P. viticola*. It remains possible that some recombination cold-spots were caused by heterozygous inversions that prevent COs in the region and thus may not be detected in a F1 progeny (Stevison et al, 2011).

The vast majority of the markers exhibited expected Mendelian segregation ratios along the entire genome. However, two regions in different LGs were affected by a significant segregation bias in the first progeny only, despite a common parent between the two crosses. Each region was affected independently in each parental map. This can be explained by a two-locus deleterious epistasis: strains Pv412 11 and Pv1419 1 present a pair of incompatible alleles on two different chromosomes. Negative epistasis between parental alleles at distinct loci is common in crosses between distinct species or subspecies, including in plant pathogens (Yuzon et al, 2023). Examples of segregation distortion due to epistatic interactions can also be found in intraspecific crosses of autogamic and allogamic plants (Törjék et al, 2006; Li et al, 2011). In Europe, *P. viticola* populations are weakly structured (Fontaine et al, 2013) and our three parent strains were collected at similar distances from each other. Thus, the incompatibility between Pv412 11 and Pv1419 1 cannot be simply explained by their distinct geographical origins.

### Crossover frequency and recombination landscape

Among the parent strains, Pv1419 1 exhibited a clearly higher genome-wide recombination rate in both male and female meiosis. Such intra-specific variation is extensively described in many species across the tree of life (Stapley et al, 2017). Variation in CO number can also be associated with variation in CO distribution (Ritz et al, 2017). In our case, the rise in recombination events in Pv1419 1 is spread across the genome. This could result from a reduced CO interference, which is tightly regulated in plants (Mercier et al, 2015) and animals (Zhang et al, 2018). A lot of species display different CO rates between male and female gametes (Lenormand and Dutheil, 2005), yet to our knowledge no data was available for oomycetes. In our *P. viticola* strains, the differences were low, albeit significant, but the trends were contradictory.

The similarity of CO distributions between parent strains points to a conserved genome architecture, with close chromosome structures and similar physical constraints during meiosis. Overall, repeat-rich and gene-poor regions were less recombinant, as observed in most eukaryotic genomes (Kent et al, 2017). Among oomycetes, a positive association between gene density and recombination was also observed in *Ph. infestans* (Matson et al, 2022).

The clustering of RXLR-like genes on a few chromosomes suggests that they evolved by the expansion of multigenic families. Evolution of effector repertoires through duplication is a common feature of oomycete genomes (Schornack et al, 2009). Paralogous effector sequences can indeed rapidly diverge to acquire new functions or escape plant recognition. Meiosis could play an important role in this process because unequal COs can generate new gene copies in effector-rich regions. Thus, frequent COs in effector-rich regions could promote rapid adaptation to host immune responses. In line with this hypothesis, recombination hotspots are enriched in secreted protein and putative effector genes in the genome of several fungal plant pathogens (Croll et al, 2015; Laurent et al, 2018; Müller et al, 2019). However, we found that these types of gene were rather under-represented in the most recombining windows of the *P. viticola* genome. Secreted protein genes are preferentially located in repeat-rich regions (Dussert et al, 2019) and we showed that these regions tend to recombine less. We hypothesize that transposable elements activity may be beneficial to enable a high turnover rate of effectors, but it could partially hinder recombination (Kent et al, 2017). In any case, virulence-related genes should not be considered to be consistently associated with highly recombining regions in oomycetes.

### Prevalence and origin of karyotypic anomalies

In our crosses, the majority of the offspring inherited a balanced set of chromosomes from both parents. Aneuploidies (2%) and triploidy (3-4%) are less prevalent than in *Ph. infestans* crosses, in which as much as 50% of the offspring carry karyotypic anomalies (Matson et al, 2022).

An open question remains how much abnormal karyotypes of *P. viticola* contribute to epidemic dynamics in the field. As it turns out, we did not identify any aneuploids in natural strains whose whole genome was sequenced so far, and only one triploid (strain Pv4168 1 from the population studied in Paineau et al (2024)). By contrast, *Ph. infestans* populations worldwide are dominated by a few pandemic clonal lineages, often polyploid (Li et al, 2017), and complex levels of aneuploidy are common. Triploidy is also found in the clonal species *P. cinnamomi* (Engelbrecht et al, 2021), and extensive aneuploidy was observed in the progeny of two clonal lineages of *P. ramorum* (Vercauteren et al, 2011).

In *Ph. infestans*, polyploid strains are thought to be fitter for asexual propagation thanks to increased heterosis, but are less fertile during sexual reproduction (Hamed and Gisi, 2013). In the case of *P. viticola*, triploid strains may be evolutionary dead-ends if they produce fewer oospores, given that they are the only overwintering form. This probably favors the maintenance of an efficient sexual reproduction that result ina majority of balanced karyotypes. Consequently, the contrasted modes of reproduction between *P. viticola* and the predominantly clonal *Ph. infestans* could explain their difference in karyotypic stability. It would be interesting to assess the prevalence of triploidy in tropical vine-growing regions where asexual propagation can take place all year round, for example in South East Brazil (Camargo et al, 2019; Santos et al, 2020).

All but one of the aneuploidies in *P. viticola* arose from uneven disjunction of chromosomes in male meiosis, in sharp contrast with *Ph. infestans* in which most aneuploidies were found to have an oogonial origin (Hamed and Gisi, 2013). One parent (Pv412 11) produced the majority of abnormal male gametes, which suggests that this tendency is subject to intra-specific variation. Strikingly, the extra chromosome sets of triploid offspring were also of paternal origin. Half of them were caused by diploid male gametes. Distinct mechanisms probably exist between female and male meiosis, which lead to a more error-prone meiosis in the latter. Our understanding of the cytological and molecular determinants of gamete formation in oomycetes remains limited, especially so when it takes place inside plant tissues. However, triploidy was also provoked by the fertilization of one oogonium by two independent haploid male nuclei (dispermy). This provides evidence that triploidy in oomycetes can arise through multiple mechanisms, not exclusively from abnormal meiosis. All three parent strains were found to cause dispermy, so it is probably relatively common in nature.

### Self-incompatibility and evolution of the MAT locus

We observed no offspring derived from selfing, confirming that sexual reproduction in *P. viticola* is strictly heterothallic. Secondary homothallism has sometimes been observed in other primarily heterothallic oomycetes, but involved either environmental stress or somatic variation disrupting the mating-type regulation system (Michelmore and Wong, 2008; Judelson, 2009).

No recombination was observed between the divergent alleles MAT-a and MAT-b of the mating-type locus, in accordance with the strong linkage disequilibrium noted by Dussert et al (2020a). Since the locus is pericentromeric, the divergence between the two alleles may have been favored by the recombination suppression in the vicinity of the centromere. This can explain the maintenance of the asymmetry of heterozygosity that determines the mating-type in this species.

The same mechanism of evolution for self-incompatibility loci has been proposed in various eukaryotic kingdoms. For example, complete linkage between the MAT locus and a centromere is observed in some ascomycete or basiodiomycete fungi (Idnurm et al, 2015). Similarly, sex-determining regions are pericentromeric in several dioecious plants (Charlesworth, 2019).

### Adaptation by Loss Of Heterozygosity

Copy-neutral LOH was observed in four individuals and affected large segments. This phenomenon has been extensively described in *Phytophthora spp* (Chamnanpunt et al, 2001; Lamour et al, 2012; Dale et al, 2019). Moreover, isolates of the downy mildew pathogen *Pe. effusa* can exhibit intermediate LOH affecting only a fraction of the nuclei (Fletcher and Michelmore, 2023). Spontaneous LOH due to mitotic recombination could also occur frequently in *P. viticola*, but may remain undetected as long as it stays limited to a minor proportion of the nuclei. In our case, LOH events must have taken place during the first step of strain propagation, as it would otherwise not have affected the entirety of the material we sequenced, and therefore not have been revealed. LOH can provide an additional way of adaptation during the growing season, through the fixation of beneficial alleles faster than annual sexual reproduction. For example, important traits such as fungicide tolerance and virulence are determined by co-dominant or recessive alleles (Lamour et al, 2012; Matson et al, 2022; Paineau et al, 2024). It has been shown that LOH can lead to a gain of virulence in *Pe. effusa* (Lyon et al, 2016).

Interestingly, two strains in the present study displayed LOH on an identical segment which corresponds to the entire long arm of the acrocentric chromosome 15. This suggests that some chromosomes may be affected by mitotic recombination more often. Similarly, LOH was observed in the same regions in different lineages of P. ramorum (Dale et al, 2019).

## Conclusion

*P. viticola* appears to be a conventional example of a highly sexual and self-incompatible organism. It istherefore an interesting plant pathogen model to study the response to new selection pressures such as the deployment of resistant varieties. As the formation of primary inoculum fully depends on sexual reproduction in temperate vine-growing regions, the recombination landscape is likely to have an impact on the efficacy of selection at different loci. The possibility of karyotypic variations should be kept in mind in future research on *P. viticola* genetics. In particular, the potential link between ploidy level and variations in fitness remains aquestion to be addressed. The linkage maps described here will be useful to guide future *de novo* chromosome-scale assemblies of *P. viticola* genomes. Moreover, they pave the way for QTL mapping of crucial traits such as virulence towards disease-resistant grapevines.

## Data availability

Allegro probe sequences, VCF files and linkage maps data are available at https://doi.org/10.57745/KX5YAQ. Whole genome Illumina DNA sequences of the parent strains are available at NCBI SRA under project number PRJNA1095879. Allegro targeted sequencing data of the parent strains and their progeny are available at NCBI SRA under project number PRJNA1107130.

## Acknowledgements

We thank Auŕelie Bérard, Isabelle Le Clainche and Damien Hinsinger (INRAE EPGV facility, National Genotyping Center, Evry, France) for carrying out Allegro Targeted Genotyping and for helpful follow-up exchanges. We acknowledge the Genotoul bioinformatics platform (Toulouse, France) for providing computing and storage resources. We thank Benôıt Laurent and Stéphanie Mariette for helpful comments on the first version of the manuscript.

## Author contributions

FD and MFO designed the study, supervised the analyses and acquired the funding. IDM and ED retrieved and propagated the grapevine downy mildew strains. ED and CC performed bioinformatic analyses. ED carried out the DNA extractions and performed genetic data analyses. ED, FD and MFO wrote the article, and all authors read and approved the submitted manuscript.

